# Speciation with gene flow via cycles of isolation and migration: Insights from multiple mangrove taxa

**DOI:** 10.1101/335166

**Authors:** Ziwen He, Xinnian Li, Ming Yang, Xinfeng Wang, Cairong Zhong, Norman C. Duke, Suhua Shi, Chung-I Wu

## Abstract

Allopatric speciation requiring an unbroken period of geographical isolation has been the standard model of neo-Darwinism. While doubts have been repeatedly raised, strict allopatry without any gene flow remains a plausible mechanism in most cases. To rigorously reject strict allopatry, genomic sequences superimposed on the geological records of a well-delineated geographical barrier will be necessary. The Strait of Malacca, narrowly connecting the Pacific and Indian Ocean coasts, serves at different times either as a geographical barrier or a conduit of gene flow for coastal/marine species. We surveyed 1,700 plants from 29 populations of five common mangrove species by large scale DNA sequencing and added several whole-genome assemblies. Speciation between the two oceans is driven by cycles of isolation and gene flow due to the fluctuations in sea level leading to the opening/closing of the Strait to ocean currents. Because the time required for speciation in mangroves is longer than the isolation phases, speciation in these mangroves has proceeded through many cycles of mixing-isolation-mixing, or MIM cycles. The MIM mechanism, by relaxing the condition of no gene flow, can promote speciation in many more geographical features than strict allopatry can. Finally, the MIM mechanism of speciation is also efficient, potentially yielding m^n^ (m>1) species after n cycles.

**Significance statement:** Mechanisms of species formation have always been a conundrum. Speciation between populations that are fully geographically isolated, or allopatric speciation, has been the standard solution in the last 50 years. Complete geographical isolation with no possibility of gene flow, however, is often untenable and is inefficient in generating the enormous biodiversity. By studying mangroves on the Indo-Malayan coasts, a global hotspot of coastal biodiversity, we were able to combine genomic data with geographical records on the Indo-Pacific barrier that separates Pacific and Indian Ocean coasts. We discovered a novel mechanism of speciation, that we call mixing-isolation-mixing (MIM) cycles. By permitting intermittent gene flow during speciation, MIM can potentially generate species at an exponential rate, thus combining speciation and biodiversity in a unified framework.

## Introduction

Speciation driven by geographical isolation with no possibility of gene flow, or strict allopatric speciation, has been the standard model of neo-Darwinism (1, 2). Although occasional exceptions are acceptable in this view (3-5), extensive violations of strict allopatry would contradict many of its central tenets. One of these tenets is the nature of species as defined by the Biological Species Concept (2). The argument for strict allopatry has usually been that gene flow would homogenize the diverging populations and retard speciation (2). After the completion of speciation, secondary contact may lead to a hybrid zone but evolutionary introgressions via extensive gene flow generally do not ensue (6).

The stringent requirement for complete geographical isolation, however, is not without difficulties. Chief among them is the paucity of geographical features that can fully stop gene flow to sustain long-term isolation. As a result, the observed biodiversity seems too extensive to rely solely on the limited opportunities for strict allopatric speciation (7). Fisher outlined a verbal theoretical model of clinal speciation (8). Endler suggested that parapatric speciation, arising between adjacent populations that continue to exchange genes at a reduced level, may be far more common than allopatry (9). Divergent selection in parapatry can be sufficient to overcome the homogenizing effects of migration if individual genic effects are examined (see ref. (10)). In this genic view, the level of divergence at the completion of speciation would be uneven across the genome. In particular, there may exist “genomic islands of speciation” (GIS) that are involved in divergent adaptation or reproductive isolation (5, 11-13).

Evidence for locus-dependent gene flow leading to the formation of GIS has been widely reported (5, 11-13). Cruickshank & Hahn rejected many reported GIS as products of processes unrelated to speciation (14). More generally, it has been pointed out that genomic data alone could not have the power to reject the allopatric model, even when GIS can be properly identified (15). In particular, if geographical isolation arises between subdivided populations, allopatry would likely be falsely rejected. Other sources of data will be necessary.

The resolution of the issue of speciation with gene flow may be possible if historical data on geographical barriers, which would offer a temporal perspective, are available. Fauna and flora of the two ocean-coasts delineated by the Indo-Pacific Barrier (IPB) are particularly suited to such inquiries (Fig. 1a). The Strait of Malacca, a main feature of IPB, can impose large-scale geographical isolation for taxa with ocean-current dependent migration. Unlike the isolation at the Isthmus of Panama, the IPB isolation is not permanent. When the sea level rose and fell periodically during the Pleistocene (16), the Strait of Malacca, which is much shallower than the two oceans, closed and opened intermittently to ocean currents and gene flow (Fig. 1b) (17). The timing of the alternation of the phases can be inferred from geological records (Fig. 1b). Hence, the DNA divergence pattern can be superimposed on the geographical records of the physical barrier itself.

**Fig. 1.**
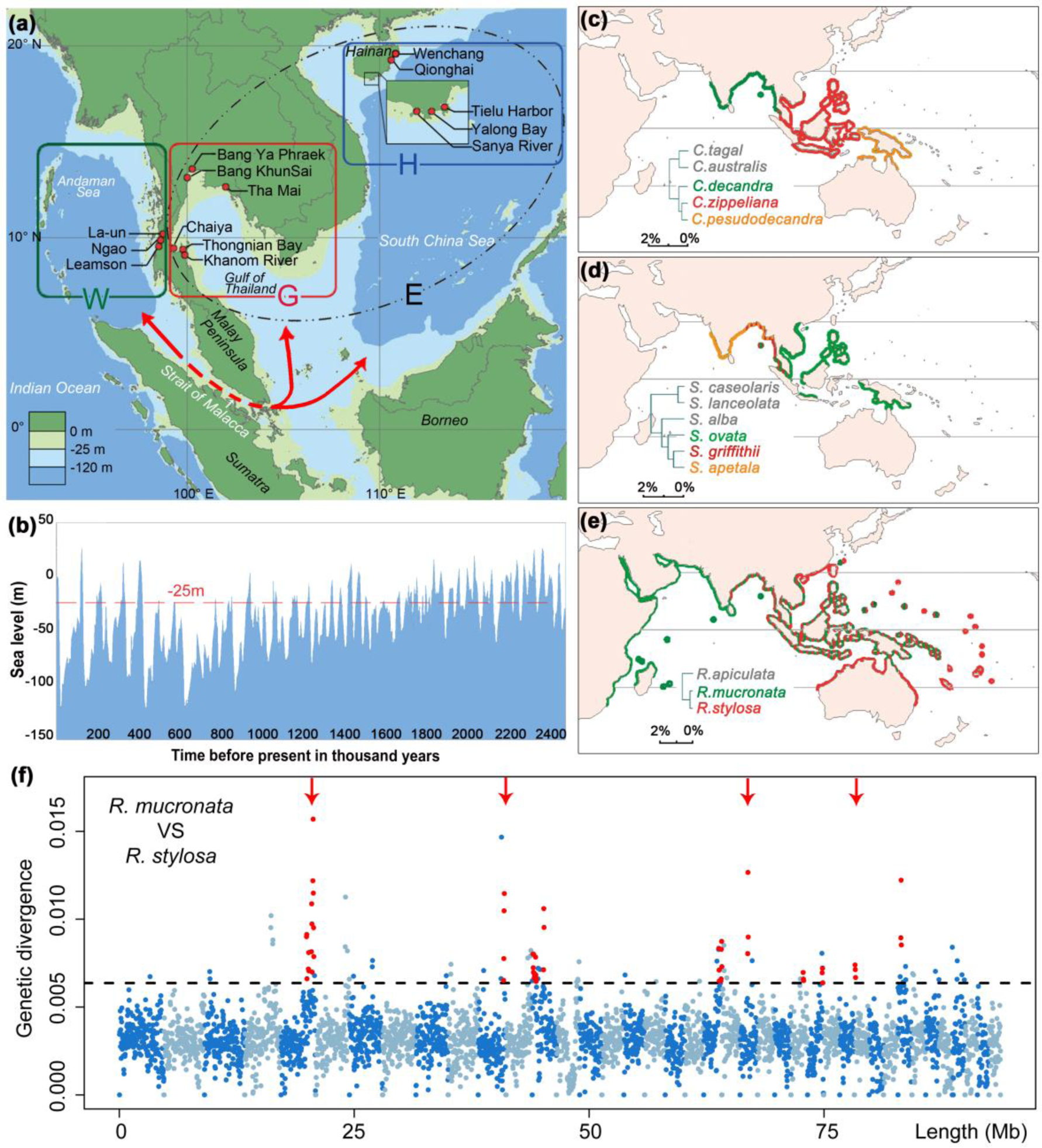
Geography and patterns of speciation in the Indo-Western Pacific. (**a**) Population samples were collected from both the Indian and Pacific Ocean coasts, separated by the Strait of Malacca. Red dots mark – sampling sites, H – Hainan Island, G – Gulf of Thailand, and W – West coast of Thailand. E stands for H + G to the east of the Strait. The red arrow-headed line depicts ocean current (and potential gene flow) through the Strait of Malacca between the Indian and Pacific coasts. (**b**) Sea level changes in the last 2.5 Myrs. The red line marks the depth of the Strait of Malacca (-25 m). (**c**) *Ceriops*; (**d**) *Sonneratia*; (**e**) *Rhizophora*. The species distribution of each genus is based on *Mangrove ID* (32). Species pairs delineated by the Strait of Malacca are shown in red and green, while the phylogeny is given in the inset. Additional species boundaries are given in Fig. S1. (**f**) Genetic divergence in 50 Kb sliding windows across the genome between *R. mucronata* and *R. stylosa*. Alternating colors denote different scaffolds; the dotted horizontal line marks the highest 5th-percentile in divergence. Red points indicate peaks of consecutive windows with elevated divergence (≥ 100 Kb). The four red arrows indicate divergence peaks that remain after controlling for mutation rate variation, as scaled by the divergence between *R. apiculata* and *R. mucronata*.

A larger issue raised by speciation mechanisms is biodiversity. There are a number of biodiversity centers globally. Among them, an exceptionally biota-rich region is found along the Indo-Malayan coasts (1, 18). Major groups of flora and fauna display unequalled species richness on these coasts (19-21). Mechanisms of speciation have been proposed, and rejected, as an explanation for exceptionally high diversity in such hot-spots (22) for two reasons. First, these centers often do not have geographical features that can facilitate allopatric speciation by imposing long-term geographical isolation (18). Second, speciation by strict allopatry (e.g., at the Isthmus of Panama) is not an efficient mechanism to generate the multitude of species because species would simply double in number. This study will attempt to connect speciation mechanisms and species richness in a single framework. Finally, given the breadth of the subject matter, necessary backgrounds and potential criticisms cannot be fully addressed in the main text. These additional topics are presented in the section *Replies to the objections to the MIM model* of Supplementary Note.

## Results

In this study, we analyze the divergence in five distantly related taxa of mangroves, which are woody plants that independently invaded the intersection between land and sea within the last 100 million years (Myrs) (20, 23-25). Because mangroves occupy a narrow band on the tropical coasts, their distributions are essentially one-dimensional, making it easier to identify geographical barriers between species. For mangroves on the two ocean-coasts (referred to as the East vs. West regions in Fig. 1a), the barrier is often the Strait of Malacca, which opens and closes periodically to ocean currents, that are conduits of mangrove seed dispersal (see Introduction). Globally, there are 70 or so mangrove species and >80% of them can be found on the Indo-Malayan coasts (26). Many of these mangrove taxa have existed and undergone diversification only in this region. In contrast, only 8 species exist in the New World tropics (20, 23, 27). Since other taxa are also highly diverse on the Indo-Malayan coasts (28, 29), the geographical mechanism of speciation in mangroves may be broadly applicable to other fauna and flora.

In this study, we approach mangrove speciation from both ends: divergence between good species and differentiation between geographical populations. By doing so, we resolve the dilemma in studying speciation. The dilemma is that good species may be too divergent to inform about speciation events (11, 14, 16, 30), but sub-species and geographical populations are not, and may not become, true species.

We have generated high-quality whole-genome sequences of multiple individuals from four species of mangroves as presented in ref. (25). For the analysis of speciation history, two genomes, one from each species, were used in this study. Many more samples but fewer genes are necessary to study population differentiation. The geographical populations of five common mangrove species on the two coasts to the East and West of the Strait are shown in Fig. 1a. Sampling was done in 14 locations of the three areas: Hainan Island in China (H), the east coast of Thailand facing the gulf (G), and the west coast of Thailand (W). In total, approximately 1,700 individuals from the five species were collected (Table S1) and subjected to sequencing. Following a published method (31), we obtained an average of 70 Kb of sequence across 80 genes per individual (Table S2).

### Speciation history in the Indo-Western Pacific (IWP)

The Indo-Malayan coasts, as part of the IWP and depicted in Fig. 1a, represent an important biodiversity hotspot. Near the tip of the Malay Peninsula, more than 20 mangrove species can be found in a local population (ref. (32, 33) and our field observations). At least nine mangrove genera had formed relatively recently originated species on these coasts and the recent speciation events (< 4% divergence) are shown in Table 1. The five genera most active in speciation during this time will be analyzed in detail here. Documented hybridizations are not uncommon in areas of sympatry (Table 1). Molecular typing has shown that hybrids are all F1 (34-39) and planting experiments have found poor pollen maturation or seed germination in the hybrids (40, 41).

**Table 1.**
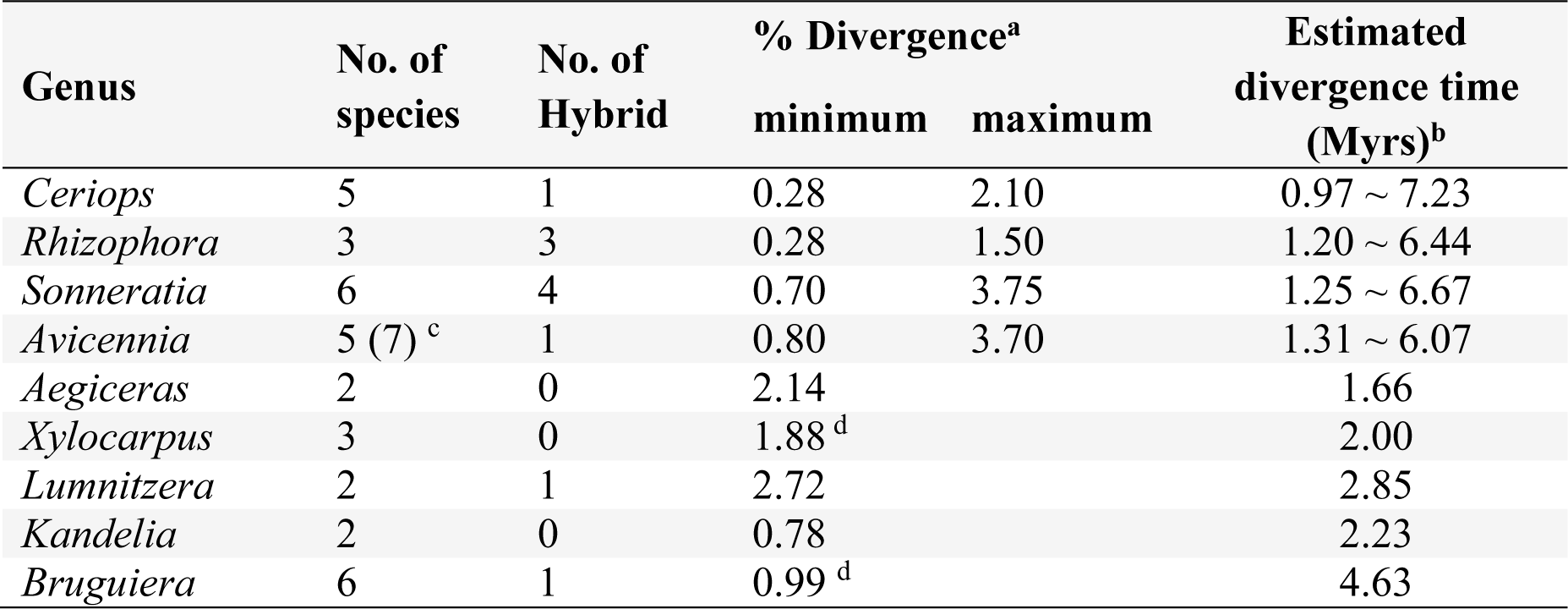
| Species Divergence within nine IWP mangrove genera

| Genus | No. of species | No. of Hybrid | % Divergence<sup>a</sup> |  | Estimated divergence time (Myrs)<sup>b</sup> |
| --- | --- | --- | --- | --- | --- |
|  |  |  | minimum | maximum |  |
| *Ceriops* | 5 | 1 | 0.28 | 2.10 | 0.97 ~ 7.23 |
| *Rhizophora* | 3 | 3 | 0.28 | 1.50 | 1.20 ~ 6.44 |
| *Sonneratia* | 6 | 4 | 0.70 | 3.75 | 1.25 ~ 6.67 |
| *Avicennia* | 5 (7)<sup>c</sup> | 1 | 0.80 | 3.70 | 1.31 ~ 6.07 |
| *Aegiceras* | 2 | 0 | 2.14 |  | 1.66 |
| *Xylocarpus* | 3 | 0 | 1.88<sup>d</sup> |  | 2.00 |
| *Lumnitzera* | 2 | 1 | 2.72 |  | 2.85 |
| *Kandelia* | 2 | 0 | 0.78 |  | 2.23 |
| *Bruguiera* | 6 | 1 | 0.99<sup>d</sup> |  | 4.63 |

The Strait of Malacca is a major geographical barrier for mangroves in the IWP (Fig. 1c-b). One example is *Ceriops decandra* vs*. C. zippeliana* (Fig. 1c), between which the extant boundary is right along the Strait. In Fig. 1d, the species boundary between *Sonneratia ovata* and *S. griffithii* is broader but also falls along the Strait. The third example is *Rhizophora mucronata* vs. *R. stylosa* (Fig. 1e). Their ranges overlap broadly on the two ocean-coasts adjacent to the Strait of Malacca but the general distributions suggest post-speciation dispersion across the Strait. *Rhizophora* is known to be better at dispersal than either *Ceriops* or *Sonneratia* (42). Three other genera are likely to have experienced post-speciation migration through the Strait of Malacca, much like *Rhizophora*. They are *Avicennia rumphiana* vs. *Av. alba, Lumnitzera littorea* vs. *L. racemosa* and *Bruguiera sexangula* vs. *B. gymnorhiza* as shown in Fig. S1. The geology of the region and the sea level records are shown in Fig. 1a-b. The East and West regions would be strongly isolated when the sea level drops below -25 meters, which is the historical norm.

It is important to point out that the Strait of Malacca connecting/separating the Pacific and Indian Ocean coasts is only one of many barriers in the IWP. Other geographical barriers can also be inferred. For example, the Torres Strait may have restricted the distributions of the sibling species *Sonneratia caseolaris* and *S. lanceolata* in northern Australia (Fig. S1a; reviewed in ref. (43)). The biodiversity in the IWP in relation to these barriers will be discussed below.

### Speciation with gene flow between the two ocean-coasts

The time of species divergence in the nine genera listed in Table 1 was estimated for each node of the phylogenetic tree based on DNA sequence data and the estimated species-specific nucleotide substitution rates (see Supplementary Note). In eight of these nine genera, the most recent species divergence time is within the last three Myrs. The oldest divergence time in Table 1 is about 6.5 Myrs ago. The most recent events within each genus generally fall in the time frame depicted in Fig. 1b, which shows the possible periods of gene flow (above -25m indicated by the red broken line).

A history of gene flow should be reflected in the genomic data because genomic segments involved in differential adaptation (in physiology, morphology, reproduction etc.) should be more divergent than the rest of the genome (10, 44). Many statistical tests have been developed to test the hypothesis by asking whether the level of divergence is “over-dispersed” across the genome. Here, we employed two methods (Fig. S2) on *R. mucronata* and *R. stylosa* (Fig. 1e), using the complete genome sequences published recently (25). In the first method, the divergence level in the genic regions is compared with that of the intergenic regions on the assumption that the former are more likely to be involved in the differential adaptation than the latter (45). The second method (46) relies on the variance in divergence across the genome. Both methods implement likelihood-ratio tests to compare the allopatric (H_0_) and speciation-with-gene flow (H_1_) models. In both methods, the null model is rejected with high confidence (P ∼ 0; Table S3), thus supporting the model of gene flow during speciation (see details for Supplementary Note). In order to identify the genomic segments most likely involved in speciation, we conducted a sliding-window analysis. Very large GIS regions between *R. mucronata* and *R. stylosa* that are unusually divergent are shown in Fig. 1f (see legends). Four of them, marked by red arrows, are more stringently called. In total, 40 GIS segments are identified for a total of 4,775 Kb, or 2.33% of the sequenced genome.

Fig. 1f follows the standard procedure in testing “speciation with gene flow” and rejects the null hypothesis. However, Yang *et al*. recently suggested that the statistical rejection is valid only for the simplest form of allopatry. For example, if gene flow occurs between geographical populations before, but not during, speciation, the null model would still be rejected, hence leading to the false rejection of allopatry (15, 47). In other words, the tests are done because the failure to reject would be biologically informative while the rejection is much less so. In cases of rejection, other types of data (e.g., geographical distributions of species and the nature of the putative barrier prior to the completion of speciation) need to be superimposed on the genomic analyses. In the remaining sections, such data will be used on geographical populations on the two ocean-coasts. The objective is to estimate the minimal time required for speciation, which will then be compared with the geological records of the geographical barrier itself.

### Differentiation between geographical populations on the two ocean-coasts

The Strait of Malacca has served as a geographical barrier leading to speciation in the past. We asked if it continues to serve as a barrier for geographical differentiation at present. Morphological observations support the inference of East-West differentiation (see Fig. 2a-b) and DNA sequence divergence provides the time depth of the geographical differentiation. The latter is usually expressed by partitioning the diversity within and between regions. Both π_R_, the genetic diversity within each area (H, G, or W), and π_T_, the total diversity of the species, are listed in Table 2 and legends, as well as Table S4 and Fig. S3. Population divergence between regions, denoted by F_ST_ = (π_T_ - π_R_)/π_T_) (48), generally follows the speciation pattern.

**Table 2.** | Differentiation rate and time between East and West populations

|  | *C. tagal* | *R. apiculata* | *S. alba* | *Av. marina* | *Ae. corniculatum* |
| --- | --- | --- | --- | --- | --- |
| **Samples and sequences** |  |  |  |  |  |
| No. of stands (H, G, W)<sup>a</sup> | 2, 1, 1 | 3, 2, 1 | 2, 2, 3 | 2, 3, 1 | 3, 1, 2 |
| Sample size (H, G, W) | 200, 100, 100 | 90, 68, 34 | 185, 100, 150 | 200, 89, 35 | 174, 50, 87 |
| No. segments (Total Kb) | 102 (76.6) | 124 (65.2) | 101 (59.6) | 150 (85.2) | 115 (57.3) |
| **Differentiation between East and West regions** |  |  |  |  |  |
| $F_{ST}$ | 0.267 | 0.48 | 0.67 | 0.72 | 0.75 |
| $(\pi_T, \pi_R)^b$ (/Kb) | (0.343, 0.251) | (1.241, 0.645) | (2.151, 0.701) | (2.519, 0.705) | (10.746, 2.640) |
| **Estimation of mutation rate and divergence** |  |  |  |  |  |
| Mutation rate, $\mu$<br>(/Kb/generation)<sup>c</sup> | $1.80 \times 10^{-5}$ | $1.63 \times 10^{-5}$ | $2.84 \times 10^{-5}$ | $3.26 \times 10^{-5}$ | $4.06 \times 10^{-5}$ |
| $N_e\mu$ in SIM (MIM) | 0.009 (0.007) | 0.029 (0.035) | 0.035 (0.030) | 0.020 (0.019) | 0.225 (0.069) |
| $N_em$ in SIM (MIM)<sup>d</sup> | 0.348 (0.420) | 0.796 (0.282) | 0.541 (0.183) | 0.265 (0.091) | 0.878 (0.135) |
| $T_{min}$ (Myrs) | 0.18 | 1.38 | 1.14 | 1.10 | 1.56 |
| $P^e$ | 0.67 | >0.999 | >0.999 | >0.999 | >0.999 |

**Fig. 2.**
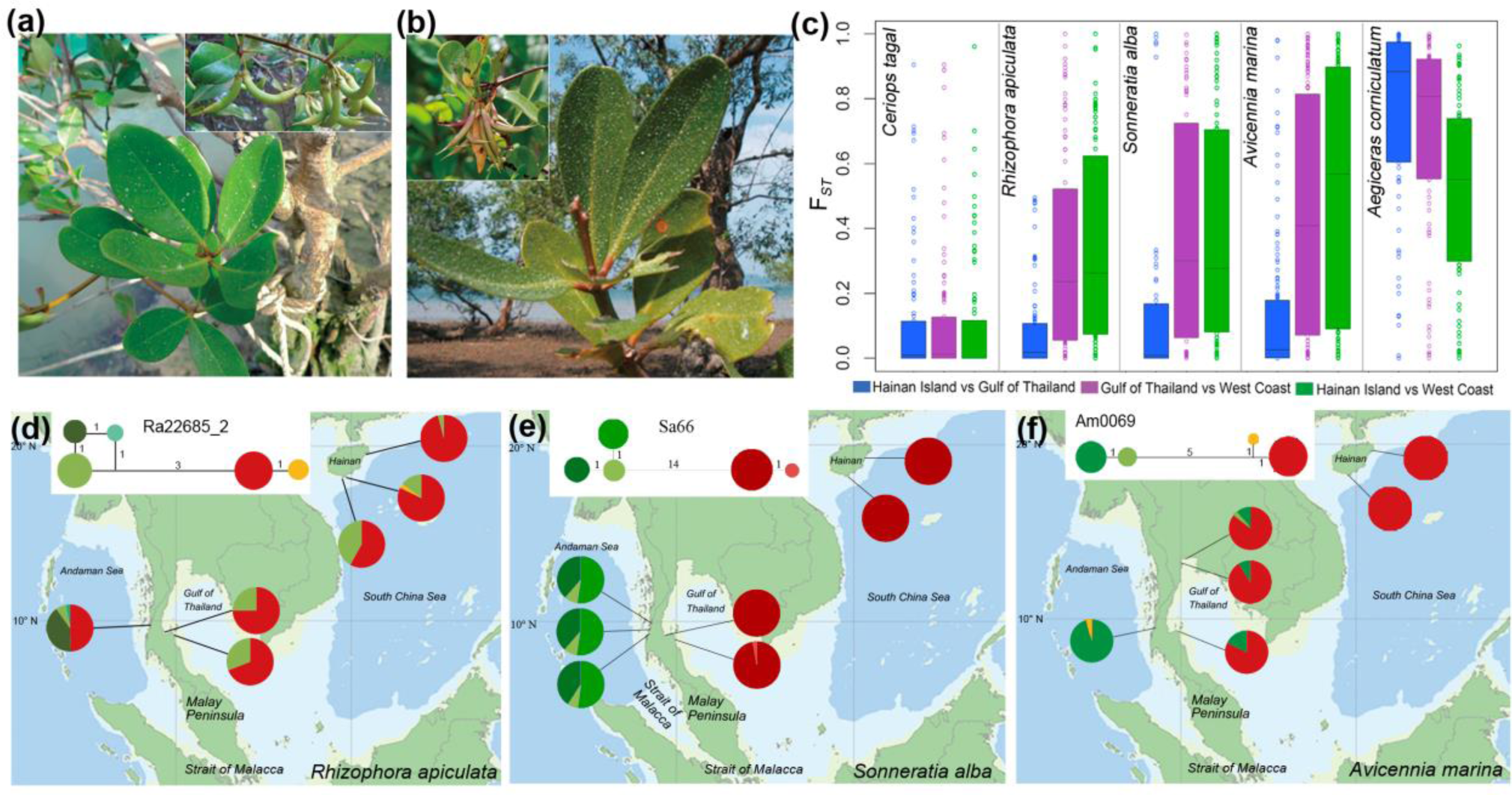
Geographical differentiation in morphology and DNA sequence. (**a** and **b**) Morphological differentiation of *Aegiceras corniculatum* from the East (**a**, samples from Hainan, China) and West (**b**. samples from Ranong, Thailand) regions. (**c**) Box plots of the F_ST_ statistic for each of the five species. For the three species with intermediate genetic diversity (*R. apiculata, S. alba* and *Av. marina*), F_ST_ between H and G is lower than the other two. (**d** to **f**) Examples of haplotype networks in the same three species that show strong East–West divergence.

One of the five species, *Ceriops tagal*, has unusually low diversity (π_T_ = 0.343 / Kb, less than 1/4 of that of the next lowest species) and hence little differentiation among all populations. Table 2 shows that all other species exhibit a larger π_T_ than π_R_ and strong population differentiation. Fig. 2c shows pairwise differentiation patterns between the three geographical areas. The divergence is relatively low in the H-G comparison in the three species with intermediate diversity (*Rhizophora apiculata*, *Sonneratia alba and Avicennia marina*), despite substantial geographical distance between the two areas. Differentiation is mainly observed between coasts of the East (combining H and G areas) and West regions (see Fig. 2c, Fig. S4). Thus, these three species suggest a key role of the Strait of Malacca in the geographical isolation between the two ocean-coasts. In the most diverse species, *Aegiceras corniculatum*, the East-West divergence is even stronger and an additional barrier (likely due to distance) also causes the divergence between the G and H populations (Fig. S5c).

Geographical differentiation can be analyzed in greater detail by analyzing haplotype structures. Three examples of haplotype networks are shown in Fig. 2d-f (see more cases in Fig. S5). The haplotypes can be clearly divided into two clades, referred to as the Eastern or Western haplotype depending on where they are more commonly found. The existence of distinct haplotypes without intermediates usually indicates strong population differentiation (49). Both the F_ST_ statistics and haplotype structures hence suggest strong differentiation between the East and West regions demarcated by the Strait of Malacca.

### DNA sequence divergence vs. geological record: How much time is needed for mangrove speciation?

Under the past sea level changes (16), the East and West regions have experienced cycles of isolation and admixture due to the repeated opening and closing of the Strait (see Fig. 1b). To compare the geological records of barrier duration with the divergence history inferred from genomic sequences, it is necessary to estimate the time required for speciation to complete (T_spp_, or speciation time). This can then be compared to the isolation time (T_iso_), the length of the periods when physical barriers to gene flow were recorded in historical data.

If we assume strict allopatry (Fig. 3a), speciation needs to be completed during geographical isolation, or T_spp_ < T_iso_. From Table 1, species divergence takes 1.2 to 6.7 Myrs with a mid-point T_spp_ of ∼ 4 Myrs. (The lowest estimate of 0.84 Myrs in *Ceriops* is less reliable due to its very low mutation rate; see Table 2). From Fig. 1b, T_iso_ is always smaller than 0.5 Myrs and rarely larger than 0.2 Myrs. Obviously, the allopatric condition of T_spp_ < T_iso_ is not met. Nevertheless, since the divergence time between good species given in Table 1 represents over-estimation of T_spp_, the rejection of T_spp_ < T_iso_ is not informative.

**Fig. 3.**
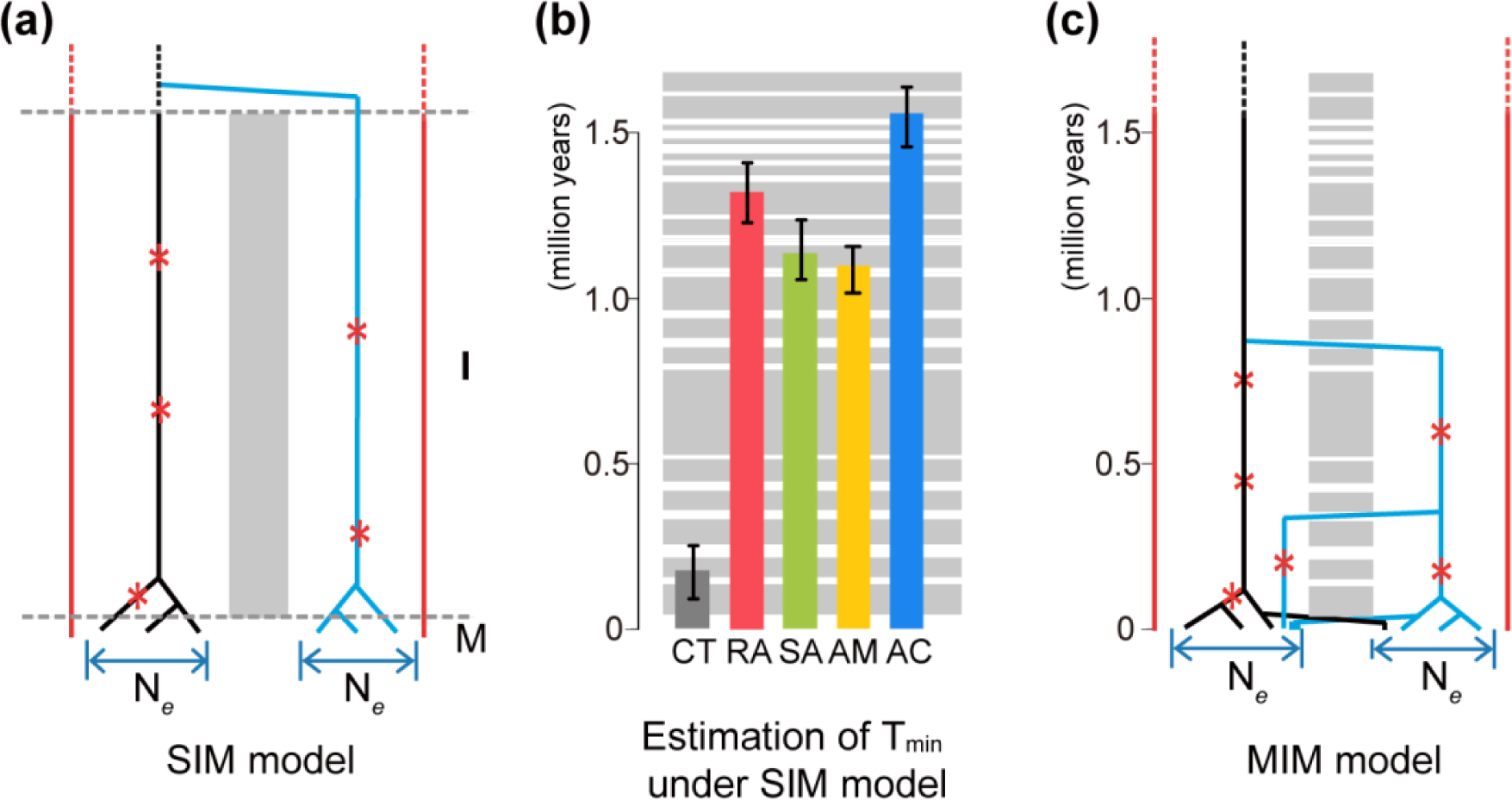
Speciation models and estimated divergence times. (**a**) The single isolation-mixing (SIM) model, equivalent to conventional allopatry. Divergence times under SIM should be relatively uniform across loci. (**b**) Estimated divergence time between the East and West populations (T_min_) under the SIM model. The shades in the background correspond to cycles of isolation and migration depicted in (**c**). Note that in four of the five species, the divergence spans multiple cycles. (**c**) The mixing-isolation-mixing (MIM) cycle model in which the cycles correspond to the geographical record of potential gene flow (Fig. 1b). Under the MIM model, the level of divergence would vary from locus to locus, depending on when migration happened.

We shall now use the lower bound estimate of T_spp_ against T_iso_. This lower bound is the divergence time between geographical populations. A new population genetic framework is developed for the purpose of estimating T_spp_ between two randomly chosen genes from the same or different populations. This new framework is presented in detail in Supplementary Note. It is distinct from previous models because it will be needed to compare the allopatric model (Fig. 3a) with our new MIM model (Fig. 3c) with multiple cycles of isolation and migration. The likelihood of observing various distributions of divergence is formulated as the function of T_spp_, N_e_ and m, where N_e_ is the effective population size, and m is the migration rate (Table 2). We then use the maximum likelihood estimates (MLE) to obtain parameters (Table 2). Note that the null model here is strict allopatry, portrayed by the single isolation-migration (SIM) cycle (Fig. 3a). If gene flow occurred during isolation, we would under-estimate T_spp_ and the rejection of allopatry would be conservative.

Fig. 3b presents the estimated T_spp_ for the five species of mangroves under the allopatric SIM model. For a comparison, the temporal sequence of migration and isolation phases at the Strait of Malacca is also shown. With the exception of *C. tagal*, the estimated T_spp_’s exceed 1.2 Myrs in the four other species. As the null model of T_spp_ < T_iso_ is rejected, T_spp_ must span several cycles of isolation-migration (see Fig. 3b).

### Speciation through MIM (mixing-isolation-mixing) cycles

Speciation in mangroves on the Pacific vs. Indian Ocean coasts had to go through cycles of isolation interspersed by episodes of gene flow, as recorded in the geological data (Fig. 3b). This mode of speciation will be referred to as the MIM cycle model. The likelihood ratio test (last row of Table 2) shows that the MIM model agrees with the observations better than the SIM model (Supplementary Note), except in *C. tagal* which has a very low species-wide polymorphism.

Under the MIM model, the distribution of neutral divergence among loci should be broader than under SIM, if everything else being equal. We use D_*max*_ (differences between the two most divergent haplotypes at any locus) as the measure. The distribution of D_*max*_ is given in Fig. 4a-c. The vertical red lines represent the average level of divergence between species or sub-species. All three species have many loci where D_*max*_ is larger than the level of (sub-)species divergence (upper panels). These loci may reflect aspects of the East-West divergence due to geographical isolation. The standard deviations of D_*max*_ are simulated and plotted (insets in Fig. 4a-c). The observations are indeed much larger than the predictions of the SIM model and fall within the simulated distributions under the MIM cycles. Thus, the divergence of mangroves on these coasts may have been influenced by periodic gene flow increasing among-locus variation.

**Fig. 4.**
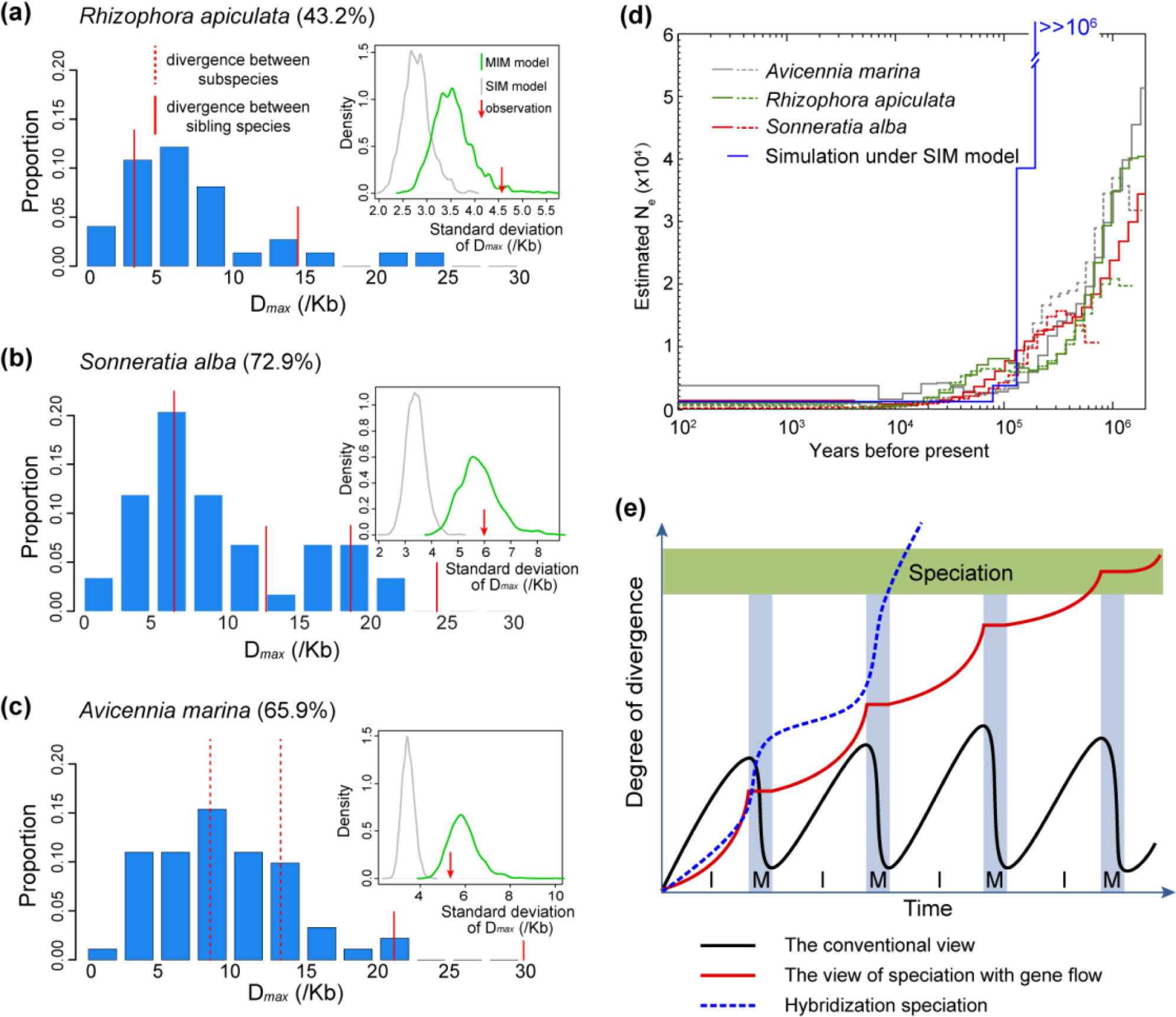
Patterns of genetic divergence under SIM vs. MIM model of speciation. (**a**) Divergence among *R. apiculata* populations. The distribution of D_max_ (differences between the two most divergent haplotypes for each gene) across loci is depicted. The bars add up to the total percentage of loci that show East-West divergence (given next to the species name). The vertical red lines indicate the level of species (solid line) and subspecies (dotted line) divergence. Note that the divergence between the geographical populations often exceeds that of subspecies, or even species. Inset figures present the standard deviations of D_max_ simulated under both MIM and SIM models. The observed value, indicated by the red arrow, is in agreement with the MIM (green line) but not with the SIM model (grey line). (**b**) *S. alba*. (**c**) *Av. marina*. (**d**) Changes in demography (population growth and differentiation) represented by the evolution of “effective population size”. The PSMC method (50), when applied to the whole genome sequencing data, can reveal changes in demography through time. Here, two individuals from each species were used, portrayed by a solid and a dotted line, respectively. Because the effective population size is sensitive to population subdivision, the analysis can discriminate between the SIM and MIM models. As shown, the population size increases gradually back in time, which is the characteristic pattern for the MIM model. In contrast, the SIM model would yield an extremely steep increase. (**e**) Three scenarios of divergence and eventual speciation. Blue shades indicate periods of migration that punctuate long periods of isolation. Speciation is indicated by high divergence. In the conventional view (black line), gene flow would reverse the divergence. Under the MIM cycles model, the level of divergence is only minimally affected by gene flow (red line). Some parameter combinations under the MIM model would underlie a third possibility (dotted blue line) whereby gene flow after a period of isolation may speed up divergence (see text).

Because isolation increases genetic variation, it also increases the effective population size. Hence, MIM and SIM models would show distinct patterns. As the genomes of three of the five species have been sequenced (ref. (25) and He *et al*., unpublished data), we re-sequenced two additional individuals for each of the three species. The PSMC method (50) infers effective population sizes at different time points in the past by comparing haploid genomes. Periods of isolation are reflected in non-coalescence and can be defined as changes in effective population size.

The PSMC results on *R. apiculata, S. alba* and *Av. marina* are given in Fig. 4d. While PSMC is usually used to model the changes in population sizes, we use it here in the context of the timing of population differentiation on the Pacific vs. Indian ocean coasts (see Supplementary Note). All three species show very small effective population sizes in the last 20,000 years, corresponding to the retreat of the last global glaciation. Going back in time, the effective population sizes increase gradually, suggesting isolated populations that have had low or intermittent gene flow during the preceding 2 Myrs. The overall PSMC patterns indicate historical gene flow spread over a long span of time, in accordance with the geological records. Had the gene flow been concentrated in a short period, the simulated SIM model would yield a steep increase in effective population size during a very short window of time (Fig. 4d).

## Discussion

Gene flow is conventionally perceived as a homogenizing force that can reverse population divergence and block speciation (black line in Fig. 4e). This has been the principal consideration of the strict allopatric model of speciation. The absence of gene flow due to geographical isolation is eventually superseded by the evolution of reproductive isolation that underpins the Biological Species Concept (2, 51). In recent years, the genic perspective suggests that gene flow during speciation would not necessarily impede divergence, as long as selection is not swamped by migration (red line in Fig. 4e) (10, 11, 52, 53). By superimposing the genomic information on the geological records, this study demonstrates that speciation on the Indo-Malayan coasts must have progressed in alternate phases of gene flow and isolation.

The MIM model therefore bridges two large sets of speciation literature. In one set, the main concern has been about the geological and phylogeographical records of speciation, which have been expertly reviewed by Hewitt (6). It is, however, not clear whether the phylogeographical literature has rejected the model of strict allopatry or has reinforced it. For example, depending on when the hybrid zone is formed, the geographical records may either suggest “speciation with gene flow”, or reinforce the view that gene flow can happen only after speciation is *fait accompli* (54, 55). In this backdrop, earlier cyclic hybridization models linking climatic cycles with speciation (56-58) are extensions of the allopatric model. In these extensions, speciation is complete in one cycle with full isolation followed by migration. The process would continue through cycles of geographical speciation and post-speciation range expansion.

A second set of the literature concerns the genomic divergence that can reveal the speciation history (10, 11, 13, 59, 60). Nevertheless, as shown by several analyses (15, 47), genomic data can inform about the occurrence of gene flow but not about when it happened. Gene flow prior to the onset of speciation might be misinterpreted to be gene flow during speciation. No less important, gene flow could be a continuous trickle or might be concentrated in short episodes of geographical panmixia, interspersed with periods of strict isolation. These isolation phases are important for the evolution of postmating reproductive isolation because incompatibility cannot evolve easily under gene flow (61, 62). In this sense, the MIM model has features of both allopatry and “speciation with gene flow”.

Interestingly, it has been posited that gene flow may even speed up speciation (the blue dotted line in Fig. 4e). This could happen if and when adaptive gene complexes, built up during isolation, are shuffled to generate many new combinations. Hybridization speciation (63-65) and adaptive radiation by hybrid swarm are such examples (66). Furthermore, many domesticated breeds were indeed created by hybridization between existing varieties (67-69). Thus, both plant and animal domestication resembles the MIM cycles, whereby breeds were separately domesticated with occasional exchange of genes. Although the idea of well-timed gene flow speeding up speciation is attractive, there is currently no evidence that it applies to mangrove speciation.

Finally, the MIM model may also bridge the gap between biodiversity and speciation studies. Many explanations have been proposed for the existence of biodiversity hotpots. Strangely, speciation has often been ruled out (70) as a mechanism of biodiversity, mainly for want of geographical features that can impose long term isolation. With MIM cycles, the stringent requirement is relaxed and many geographical features could conceivably drive speciation. In the IWP, because the sea floor in the Indo-Malayan region has been relatively high, many shallow barriers have existed throughout the region (71). When the global sea level began to decrease and fluctuate around that lower level 25 Myrs ago (16), many parts of the Indo-Malayan coasts may have experienced cycles of isolation and admixture. Indeed, as Renema *et al*. have pointed out, species diversity in the Indo-Malesia started to increase during Miocene (22, 72).

The MIM cycle mechanism may be applicable to other high-diversity spots as well. In the same time frame as mangrove speciation on the Indo-Malayan coasts, islands of the Aegean Archipelago in the Mediterranean may have been repeatedly connected and disconnected due to sea level changes. The radiation of the annual plants in the genus Nigella across the archipelago (73) could also be driven by a mechanism like MIM. In a most dramatic setting, Lake Victoria, which has experienced repeated rises and falls of water level, harbors an extraordinary diversity of cichlid fish (74). Diverse flora in neo-tropical rain forests has also been attributed to periods of cooler and drier climates driven by the cyclical glacial events (75). In addition, fragmentation and reconnection of high elevation habitats during the late Pleistocene has been proposed as an explanation for avian diversification in the neotropics (76).

When diverging populations become full-fledged species, migration in the next M phase would be equivalent to range expansion. If speciation occurs after each isolation phase, there can be as many as 2^n^ species after n cycles (56). In that sense, the migration phase in the MIM cycles would play an added role in the evolution of biodiversity. More generally, isolation may create i fragmented populations. If speciation is achieved after j cycles, then the number of species after n cycles would be [i]^n/j^. In other words, the number of species after n cycles can potentially be m^n^ where m = i^1/j^ > 1. In the special case of i = 2 and j =1, m = 2^n^. Centers of high biodiversity are fascinating phenomena with many possible causes (19-21, 77, 78). We suggest that efficient speciation mechanisms like MIM cycles may play a role.

## Materials and Methods

### Geographical distribution of mangrove species in IWP

The geographical distribution of each of the nine mangrove genera in the IWP (*Kandelia*, *Aegiceras*, *Lumnitzera*, *Ceriops*, *Xylocarpus*, *Bruguiera*, *Avicennia*, *Rhizophora* and *Sonneratia*) was compiled from *World Mangrove ID* (32). Species distribution ranges of *Ceriops* were updated according to Tsai *et al*. (*79*). The distributions of *Rhizophora* and *Sonneratia* in China were updated from the field survey data of Wang and Chen (80).

### Scanning the genome for speciation islands

To identify genomic regions highly divergent between *R. mucronata* and *R. stylosa*, we performed a genome-wide divergence scan using absolute measures of differentiation. Re-sequencing data of one *R. mucronata* individual from Ranong, Thailand, and one *R. stylosa* individual from Hainan, China, were generated using Illumina HiSeq 2000 platform. Reads were mapped to the *R. apiculata* reference genome using the BWA software (81). Heterozygous sites were called using the GATK pipeline (82). We used sliding windows to scan divergence levels between the two species. We set the window size to 50 Kb and step size to 25 Kb. Windows with fewer than 10,000 sequenced sites were discarded. Divergence level of each retained window was calculated as the number of differentiated sites divided by the number of sequenced sites. Divergent sites were defined as loci homozygous within each species but different between taxa.

### Sampling, sequencing, and mapping

We collected leaf material from populations of five mangrove species from 15 stands in the three regions as shown in Fig. 1a and Table S1. For each species, at least one stand was sampled in each region and 19 to 100 individuals were collected from each stand. Intervals between sampled individuals were at least five meters. Sequencing protocols were as described in our earlier work (31). Equal amount of leaf material from each individual in every stand was mixed before DNA extraction. Based on sequences from cDNA libraries of the species, we designed primers for over 150 loci for each species. We succeeded in amplifying approximately 70 genes per species (Table S2) and sequenced them using the Illumina GA-II/HiSeq 2000 platform. The short reads sequence data were deposited in NCBI, BioProject: PRJNA303892. We mapped short reads to references using MAQ (83) with main parameters -m 0.002 and -e 200 and the parameter -q 30 to filter low-quality reads. To reduce sequencing errors, we ignored bases that were (1) located in the first 11 bp or the last 7 bp of the mapped reads, (2) with base quality less than 22, and (3) with minimum coverage less than 100. Putative single nucleotide polymorphisms (SNPs) were called if the minor allele frequency was > 0.01.

### Haplotype inference

Using the linkage information for SNPs in each pair of short reads, we estimated haplotypes and their frequencies using an expectation-maximization algorithm (84, 85). We divided the genes into two or more segments if the distance between two SNPs was longer than the length covered by paired reads. To validate the estimated haplotype phases, we sequenced 360 alleles in eight populations using the Sanger method (Supplementary Dataset 1). Haplotypes and their frequencies estimated using these two approaches were very similar. Short reads were informative for our haplotype analyses thanks to large sample sizes. We constructed haplotype networks for each gene segment based on the inferred haplotypes.

### Estimating nucleotide diversity and population structure

Using the obtained haplotype profiles, we estimated nucleotide diversity (π, the average number of nucleotide mismatches per site between two sequences (86)) within a stand/population (π_s_), region (π_R_, with all areas weighted equally), and species (π_T_, with all three regions weighted equally) for each haplotype segment. We employed F-statistics at different levels to measure population differentiation (*F*_*ST*_ = 1 – *π* _*S*_ / *π*_*T*_) (48).

After carefully reviewing haplotype networks for the five species, we observed that haplotypes of many genes could be clustered into two distinct clades corresponding to the samples from the East and West Indo-Malayan regions. We therefore clustered the haplotypes into two clades using complete linkage method hierarchical clustering analysis (84). We included segments with more than two SNPs, or two SNPs and two haplotypes. We calculated frequencies of haplotype clades in both regions.

### Demographic models and parameter estimation

We used a maximum likelihood method to estimate effective population size (N_e_) and migration rate (m) for the SIM and MIM models. The SIM model requires an additional parameter, the isolation time imposed by the geographical isolation, as depicted in Fig. 3a. The time elements in the MIM model were defined by the geological records of sea level changes. The mutation rate was inferred from exome/transcriptome data with the fossil records as described in the Table 1 legend.

The number of divergent nucleotides between two sequences sampled from the same populations was denoted as *D*_*w*_, while differentiation between populations was denoted as *Db* (w stands for within population and b for between populations). The log-likelihood function can be constructed as follows:

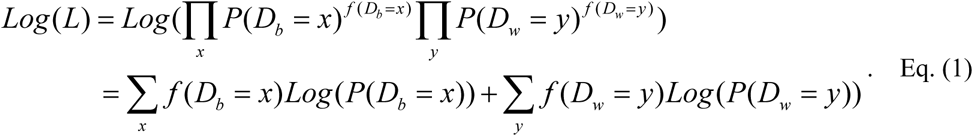

*f* (*D*_*b*_ = *x*) is the observed number of sequence pairs between populations where *D*_*b*_ is equal to x and *f* (*D*_*w*_ = *y*) is the observed number of sequence pairs within a population where *D*_*w*_ is equal to y. The probability P of *D*_*b*_ and *D*_*w*_ could be deduced using the transition probability matrix during the M phase and I phase, according to the coalescent process. The detail equations are given in the Eq. S2-S9 in Supplementary Note.

We wrote *Mathematica* scripts to obtain maximum likelihood estimates of effective population size and migration rate for the MIM and SIM model using numerical methods. Given a generation time equal to 20 years, the MIM model parameters j and k were set to 5,000 and 500 generations for each I and M phase according to geographical evidence of the recent cycles. In SIM, j is the additional parameter to be estimated.

To validate method accuracy, we carried out a series of simulations. We used the ms (87) to simulate sequences under the MIM and SIM models for 1,000 replicates for each set of parameters (Fig. S6). When isolation time was set to 1 Myrs, the standard deviation for 1,000 simulation results was no more than 0.1 Myrs. The estimation under MIM was also comparably accurate (Fig. S7).

### Simulations of DNA sequence evolution

We used ms (87) to simulate sequence evolution under the MIM scheme for 2,000 replicates in each species. Parameter values used in simulations are listed in Table 2. Six statistics were obtained from the simulated sequences for each species: D_*within*_ (average divergence within region), D_*between*_ (average divergence between regions), D_*clade*_ (differentiation among the most recent common ancestors of each clade), D_max_ (differences between two most divergent haplotypes), P_*total*_ (total number of SNPs), and F_ST_ among regions. The simulated distributions of the six statistics are comparable to values observed from data (Figs. S8-S12).

We also simulated 2,000 replicates under the SIM evolution scheme using the parameters listed in Table 2 for calculating D_*max*_. We calculated the standard deviation of D_*max*_ among genes in each relicate derived from the SIM or MIM model. The distributions of the 2,000 standard deviations from the two models are depicted in the insets of Fig. 4a-c.

To test whether the MIM model fits observed data better than the SIM model, we obtained maximum likelihood estimates of the two models for sequences simulated under SIM model. We calculated the differences in the likelihood values (Diff = log-likelihood of the MIM model – log-likelihood of SIM model) for each of the 2,000 repetitions. For *R. apiculata*, *S. alba*, *Av. marina* and *Ae. corniculatum*, the Diff value of the real data is larger than all the Diff values of the simulated sequences. Hence, the probability that the SIM model fits data better than the MIM model is less than 0.001. For *C. tagal*, the probability is 0.33. As discussed in the main text, the unusually low genetic diversity of *C. tagal* makes it powerless to compare models.

### Estimating effective population size change using whole-genome sequence data

To estimate past effective population size, we used the pairwise sequentially Markovian coalescent analysis (PSMC) (50). We used the whole-genome sequence data from six individuals (data deposited in NCBI, BioProject: PRJNA298659). *Av. marina* and *S. alba* individuals were from the Gulf of Thailand and the West Coast. *R. apiculata* samples were from Sanya and Wenchang. We mapped the resequencing data generated by Illumina HiSeq 2000 platform to the corresponding draft genomes (ref. (25) and He *et al*., unpublished data) using BWA (81). The parameters of PSMC estimation were: - N25 -t15 -r5 -p “4+25*2+4+6”. Generation time was set to 20 years. The mutation rate for each species is given in Table 2.

We also produced simulated sequence data for PSMC analysis (see Fig. 4d and Figs. S13-S15). The simulated sequences were generated by msHOT (88) with the following parameters: mutation rate (μ) set as 0.5, 1.0, 2.0 ×10 ^-9^ /site/year, migration rate (m) as 1×10 ^-4^, 5×10 ^-4^, 10×10 ^-4^ per generation, population size (N) as 100, 500, 1,000 and 5,000. Each simulated genome contained 500 loci and the length of each gene was set to 200 Kb. The recombination rate was set to 1×10 ^-9^ /site/generation.

## Acknowledgements

We thank David Jablonski, Roger Butlin, Richard Abbott, Trevor Price, Loren Rieseberg, Nick Barton, Patrik Nosil, James Mallet, Daven Presgraves, Dolph Schluter, Fangliang He, Chuck Canon and Xionglei He for comments that improved the manuscript. This study was supported by the National Natural Science Foundation of China (91731301, 91331202 and 31600182); the National Key Research and Development Plan (2017FY100705); the 985 Project (33000-18821105) and the Fundamental Research Funds for the Central Universities (17lgpy99).

## Database deposition

Illumina reads are available at the Short Read Archive under the NCBI BioProject PRJNA303892 and PRJNA298659.

